# Exploring and analysing immune single cell multi-omics data with VDJView

**DOI:** 10.1101/613083

**Authors:** Jerome Samir, Simone Rizzetto, Money Gupta, Fabio Luciani

## Abstract

**Background:** Single cell RNA sequencing provides unprecedented opportunity to simultaneously explore the transcriptomic and immune receptor diversity of T and B cells. However, there are limited tools available that simultaneously analyse large multi-omics datasets integrated with metadata such as patient and clinical information.

**Results:** We developed VDJView, which permits the simultaneous or independent analysis and visualisation of gene expression, immune receptors, and clinical metadata of both T and B cells. This tool is implemented as an easy-to-use R shiny web-application, which integrates numerous gene expression and TCR analysis tools, and accepts data from plate-based sorted or high-throughput single cell platforms. We utilised VDJView to analyse several 10X scRNA-seq datasets, including a recent dataset of 150,000 CD8+ T cells with available gene expression, TCR sequences, quantification of 15 surface proteins, and 44 antigen specificities (across viruses, cancer, and self-antigens). We performed quality control, filtering of tetramer non-specific cells, clustering, random sampling and hypothesis testing to discover antigen specific gene signatures which were associated with immune cell differentiation states and clonal expansion across the pathogen specific T cells. We also analysed 563 single cells (plate-based sorted) obtained from 11 subjects, revealing clonally expanded T and B cells across primary cancer tissues and metastatic lymph-node. These immune cells clustered with distinct gene signatures according to the breast cancer molecular subtype. VDJView has been tested in lab meetings and peer-to-peer discussions, showing effective data generation and discussion without the need to consult bioinformaticians.

**Conclusions:** VDJView enables researchers without profound bioinformatics skills to analyse immune scRNA-seq data, integrating and visualising this with clonality and metadata profiles, thus accelerating the process of hypothesis testing, data interpretation and discovery of cellular heterogeneity. VDJView is freely available at https://bitbucket.org/kirbyvisp/vdjview.

## Background

Immunological studies have revealed a surprisingly high level of heterogeneity between immune cells, even in those with same clonotype and surface phenotype, suggesting that lymphocyte populations of apparently similar phenotype could have different functions [1]. With the advent of single cell RNA-sequencing (scRNA-seq), it is now possible to unravel the heterogeneity of T and B cells and link receptor clonotype diversity to the gene expression profile of each cell and to clinical or other metadata. Multi-modality single cell datasets are rapidly pervading in medical research, and are being used to identify novel cellular states and molecular features of diseases [2-4], to extract information on the DNA (mutations, methylation), mRNA (gene expression profiles) and to further study the heterogeneity of immune cells of apparently similar clonotype and phenotype [3].

With the recent availability of scRNA-seq derived clonal and transcriptomic data, several software packages have been developed for the downstream analyses of these data types [3]. For instance software packages such as TRACER [5] BRACER [4] and VDJPuzzle (for both TCR [6] and BCR [2]) can accurately identify the full-length TCR and BCR from the sequenced cDNA. A vast set of tools are already available to perform gene expression analysis, including clustering, differential expression, dimensionality reduction, trajectory inference, and gene signature identification (e.g. https://www.scrna-tools.org/). More recently, epitope barcoding on cell surface has been also integrated with scRNA-seq, further highlighting the importance of multi-modal single cell technologies [7, 8].

Integrating these levels of genomic information can be important to fully decipher the changes of immune cells during immune response, or to identify subsets of rare cells with specific phenotypes. Tools that integrate several of the available methods to analyse single cell transcriptomics have been proposed [9, 10]. Additionally, it is often necessary to link this information with clinical and other metadata, for instance with the tissue origin, surface phenotype (e.g. flow cytometry data at the time of index sorting), or with the sample origin and disease diagnosed. To date, there are limited software packages that are accessible to non-bioinformatics experts, and that allow simultaneous analysis of gene expression, immune receptors and notably clinical and other metadata. For instance, Loupe Cell Browser 3.1 from 10X Genomics provides users with a first line of analysis to explore gene expression and annotate their dimensionality reduction plots with immune receptor information. However, such tools do not permit extensive analysis of the data, such as hypothesis testing and integration of metadata into differential expression or immune receptor analyses. Additionally, such tools usually have strict input requirements, with Loupe Cell Browser 3.1 not allowing users to analyse datasets from different technologies, such as plate-based sorting, which remains a common technology of choice to study immune repertoires.

Multi-layer analyses often require lengthy integration of bioinformatics and biological skills. Experience with software tools, such as R, is often a barrier to entry, with most of the data manipulation, visualisation and package integration being left to the user. To properly answer and address biological questions, multiple packages need to be complemented with ad hoc scripts that modify input data, filter cells and then test hypotheses, which is a source of latency between the biologist and the bioinformatician. Here, we report a novel bioinformatics tool, VDJView, which delivers an integrated set of novel and publicly available tools to analyse and visualise the clonal and transcriptomic data with clinical and metadata. The shiny app, VDJView, addresses the drawbacks in currently available multi-omics analysis tools, by removing the need for a skilled bioinformaticians, allowing researchers to test hypotheses and explore the relationship between multi-modal single cell datasets.

## Implementation

VDJView is an R Shiny web-application developed for the analysis of clonal and transcriptomic single-cell data (Figure 1). The intuitive graphical user interface allows researchers with or without computational training to interactively analyse and explore their datasets, interrogating the results against user uploaded cell metadata. VDJView acts as a wrapper for commonly used transcriptomic and receptor analysis packages (Table 1), integrating them and allowing the user to generate and manipulate figures and tables. The plots generated are exportable to publication-quality pdf files, and all tables can be downloaded in CSV format. VDJView has been extensively tested on Linux and MacOS, with most features functional on Windows as well, and has the sole requirement of an R version of at least 3.4.1 being installed. VDJView has been tested on multiple datasets available from published literature using SmartSeq2 and 10X libraries (see below). The performance of the application is detailed in Supplementary Note 1.

**Table 1:**
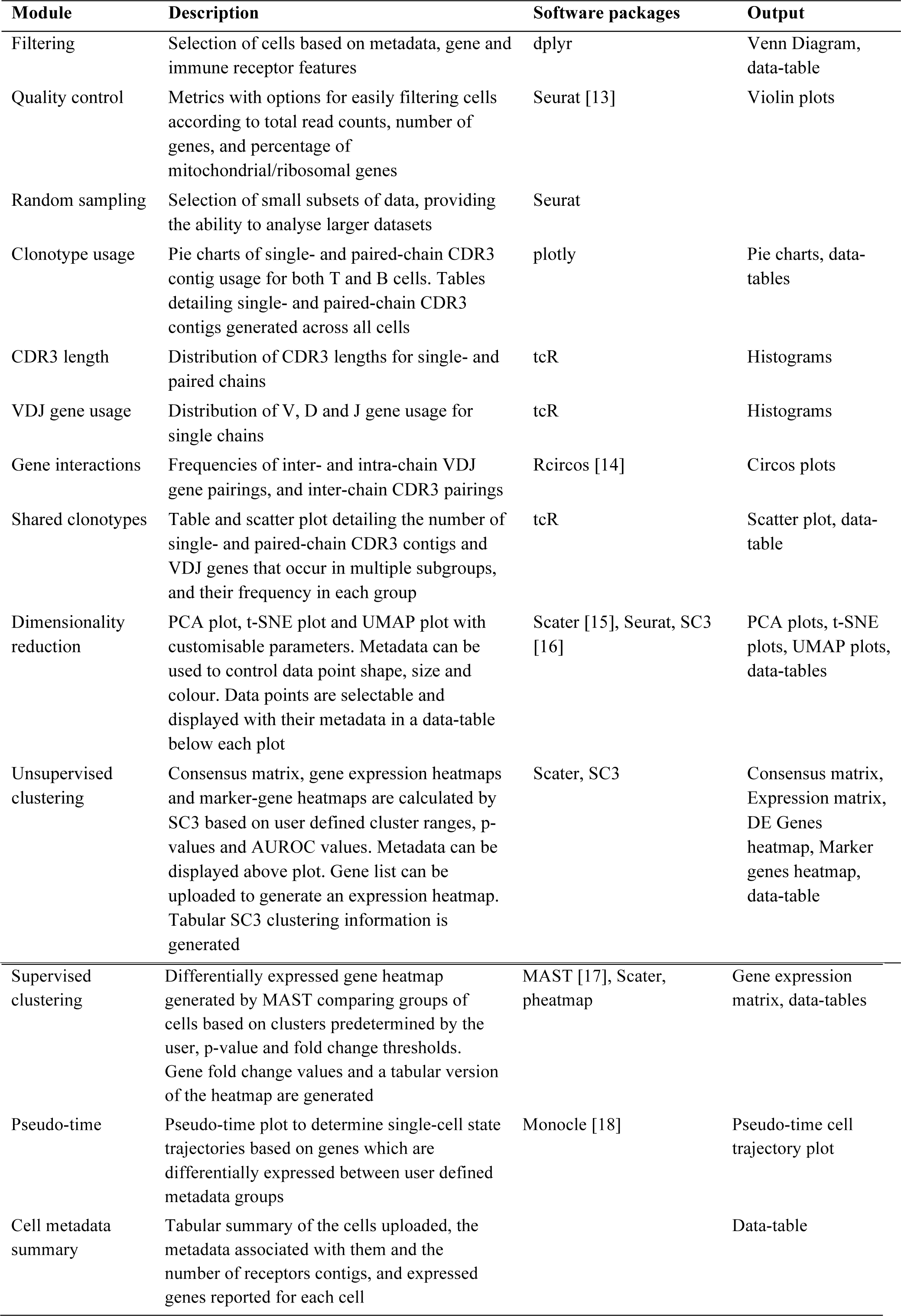
List of modules implemented in VDJView with their outputs and integrated packages.

| Module | Description | Software packages | Output |
| --- | --- | --- | --- |
| Filtering | Selection of cells based on metadata, gene and immune receptor features | dplyr | Venn Diagram, data-table |
| Quality control | Metrics with options for easily filtering cells according to total read counts, number of genes, and percentage of mitochondrial/ribosomal genes | Seurat [13] | Violin plots |
| Random sampling | Selection of small subsets of data, providing the ability to analyse larger datasets | Seurat |  |
| Clonotype usage | Pie charts of single- and paired-chain CDR3 contig usage for both T and B cells. Tables detailing single- and paired-chain CDR3 contigs generated across all cells | plotly | Pie charts, data-tables |
| CDR3 length | Distribution of CDR3 lengths for single- and paired chains | tcR | Histograms |
| VDJ gene usage | Distribution of V, D and J gene usage for single chains | tcR | Histograms |
| Gene interactions | Frequencies of inter- and intra-chain VDJ gene pairings, and inter-chain CDR3 pairings | Rcircos [14] | Circos plots |
| Shared clonotypes | Table and scatter plot detailing the number of single- and paired-chain CDR3 contigs and VDJ genes that occur in multiple subgroups, and their frequency in each group | tcR | Scatter plot, data-table |
| Dimensionality reduction | PCA plot, t-SNE plot and UMAP plot with customisable parameters. Metadata can be used to control data point shape, size and colour. Data points are selectable and displayed with their metadata in a data-table below each plot | Scater [15], Seurat, SC3 [16] | PCA plots, t-SNE plots, UMAP plots, data-tables |
| Unsupervised clustering | Consensus matrix, gene expression heatmaps and marker-gene heatmaps are calculated by SC3 based on user defined cluster ranges, p-values and AUROC values. Metadata can be displayed above plot. Gene list can be uploaded to generate an expression heatmap. Tabular SC3 clustering information is generated | Scater, SC3 | Consensus matrix, Expression matrix, DE Genes heatmap, Marker genes heatmap, data-table |
| Supervised clustering | Differentially expressed gene heatmap generated by MAST comparing groups of cells based on clusters predetermined by the user, p-value and fold change thresholds. Gene fold change values and a tabular version of the heatmap are generated | MAST [17], Scater, pheatmap | Gene expression matrix, data-tables |
| Pseudo-time | Pseudo-time plot to determine single-cell state trajectories based on genes which are differentially expressed between user defined metadata groups | Monocle [18] | Pseudo-time cell trajectory plot |
| Cell metadata summary | Tabular summary of the cells uploaded, the metadata associated with them and the number of receptors contigs, and expressed genes reported for each cell |  | Data-table |

**Figure 1.**
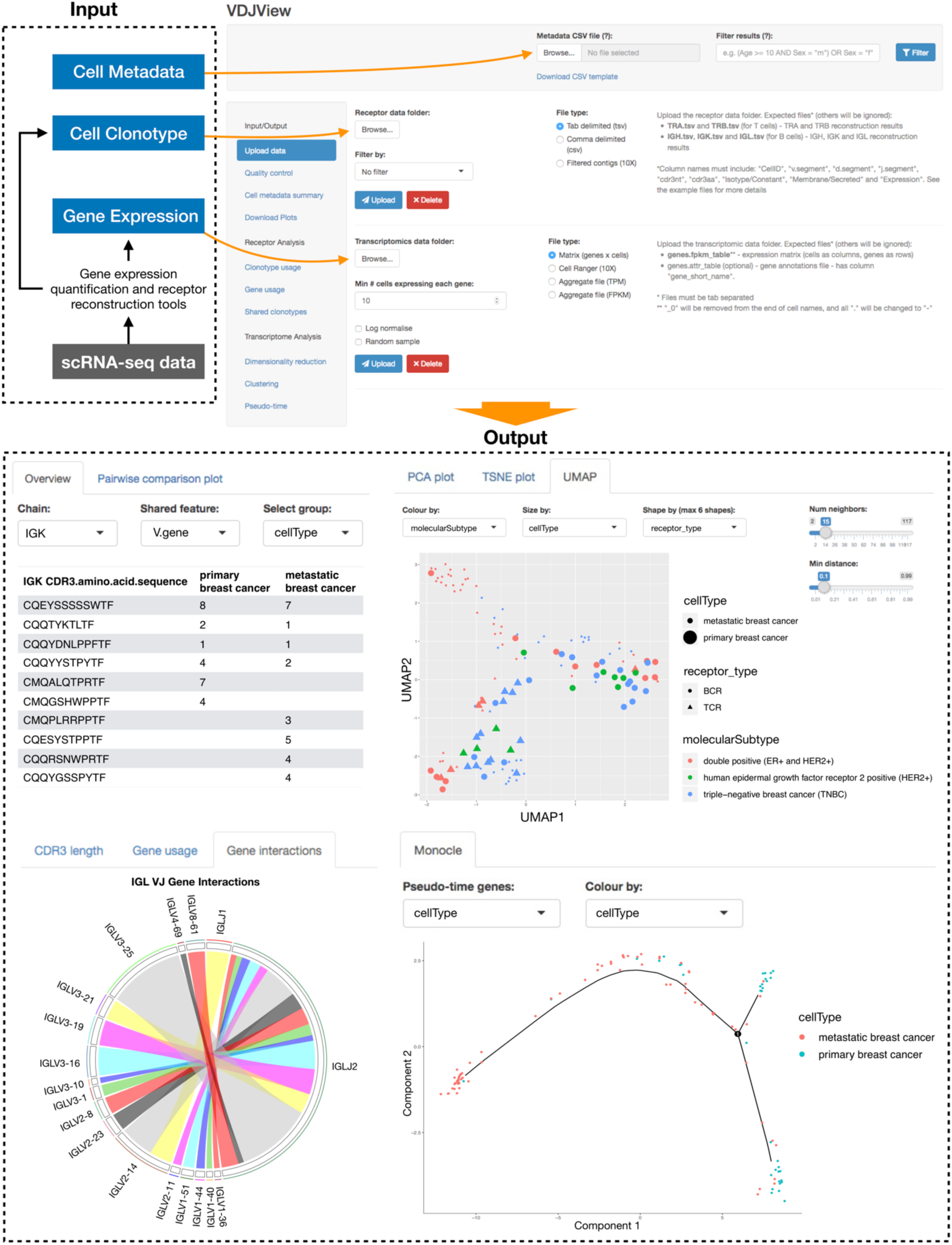
Overview of VDJView. **Top**: VDJView upload page, showing where required (immune receptor sequences and gene expression matrix) and optional inputs (metadata) can be uploaded. **Bottom**: examples of analysis using scRNA-seq from primary cancer tissues and metastatic lymph-node revealing clonally expanded T and B cells. The table (top left) shows a clonal expansion of IGL chains across primary breast tissue and metastatic lymph-node. The Circos plot (bottom left) shows the IgL V and J gene pairings identified. Dimensionality reduction using UMAP (top right) shows a cluster of B cells derived from metastatic lymph-node in two patients with ER^+^ HER2^+^ breast cancer, while T and B cells from the primary breast cancer tissue had similar gene signature regardless of molecular subtype. Pseudo-time plot (bottom right) shows the inferred evolutionary trajectory between all immune cells determined by genes that differentiate primary from metastatic tissues in two subjects with matched samples.

### VDJView input data

Pre-analysed scRNA-seq data can be directly uploaded into VDJView. The three data types that VDJView accepts are; T and/or B cell receptor data, gene expression data, and metadata. Immune receptor data can be uploaded as a list in csv or other tabular formats. Gene expression data can be uploaded as a matrix of expression counts per cell or other common formats including those generated by the 10X Cell Ranger kit. Metadata can be uploaded in csv format. VDJView can also be pipelined with computational tools that generate gene expression and immune receptor sequencing from raw data, allowing users to use their own workflow. Here, we have tested VDJView with scRNA-seq data available publicly and generated by high-throughput 3’ or 5’ end technologies, 10X and SmartSeq2 data.

### Datasets analysed

1. SmartSeq2 breast cancer T and B cells, N = ∼560 [11]
2. 10X CD8+ T cells, N = ∼150,000 (https://www.10xgenomics.com/resources/application-notes/a-new-way-of-exploring-immunity-linking-highly-multiplexed-antigen-recognition-to-immune-repertoire-and-phenotype/)
3. 10X non-small lung cancer cells (NSCLC), N = ∼8,300 (https://support.10xgenomics.com/single-cell-vdj/datasets/2.2.0/vdj_v1_hs_nsclc_5gex)
4. Hepatocellular carcinoma T cells, N = ∼4800 [12]

### VDJView features and modules

VDJView integrates multiple R software packages to provide a powerful yet cohesive repertoire of analysis modules (Table 1). Numerous interactive and customizable figures are provided for the analysis of clonotype data, and further modules are available for the simultaneous or isolated exploration of expression data. All figures and tables are updated automatically if any of the relevant parameters are changed during the analysis. Further details and a complete list of features can be found in Supplementary Note 1.

## Results

### Analysis of SmartSeq2 breast cancer cells

To demonstrate the utility and novelty of VDJView, we analysed scRNA-seq data (full-length transcriptome, SmartSeq2 protocol) from the primary breast tissues and metastatic lymph nodes of 11 subjects [11]. We applied VDJPuzzle [2] to the scRNA-seq data to quantify the gene expression and reconstruct the receptors of the T and B cells, inputting the results into VDJView. We found 170 single B cells with at least one full-length H, L or K chain, of which 101 had a full-length heavy and light chain. Similarly, we found 42 single T cells with at least one full-length α or β TCR chain, of which 30 had paired TRα and TRβ chains. Thus, we have uniquely identified T and B cells via their receptor, and confirmed the findings achieved by the authors of the original work by identifying immune cells through gene enrichment analysis [11]. In addition to these, we found 33 cells with TCR and BCR chains, suggesting that they were likely contaminants or doublets. While our analysis of the T cells revealed a highly diverse repertoire (Supplementary Figure 1), we identified a clone in BC03 which was present in both primary and metastatic lymph node tissues, as well as 31 B-cell clones, with clonotypes shared across primary and metastatic tissues, and across subjects (Figure 1 and Supplementary Figures 1-2, Supplementary Tables 1-2). Pseudo-time analysis on these immune cells showed that a common repertoire of B cells is involved in breast cancer with a migratory pattern between primary and metastatic tissues (Figure 1). We used VDJView to integrate immune receptor information with the gene expression profile and available metadata, and performed unsupervised clustering. The unsupervised clustering (Supplementary Figure 4) revealed evidence of 8 clusters based on identity (B and T cells), B-cell isotype, tissue of origin and cancer molecular subtype. T cells largely formed a single cluster with marker gene CD96 associated to immune modulation, as well as expression of IL2R-γ and FYB which is known to control IL-2 secretion. The remaining clusters were largely composed of B cells based on tissue of origin, molecular subtype of cancer, and notably a cluster that was composed of IgG1 B cells in metastatic lymph-node of double positive breast cancer, expressing gene signature suggesting they are highly active and differentiated B cells, e.g. plasmablast following a reactivation of memory B cells. In this cluster, the over-expression of PAX5 and TCL1A could also indicate presence of malignant immune cells as these genes are often found in leukemia and likely to contribute to oncogenesis BCL6 [19, 20]. Further analysis of this data is detailed in Supplementary Note 2.

### Analysis of 10X antigen specific CD8+ T cells

To further demonstrate the utility of VDJView, we have analysed the recently published scRNA-seq data with TotalSeq and dextramer stained CD8+ T cells. This dataset contains single cell data on over 150,000 CD8+ T cells isolated from 4 healthy donors, two of which were CMV positive, 44 dextramers were simultaneously used in each subject to isolate antigen specific T cells across viral infections (CMV, EBV, HPV, Influenza, HIV), and cancer (e.g., MART, MAGE NY-ESO). We used this data to study the clonal distribution within and across specific antigens and link this information to the gene expression and other metadata.

In this analysis, we uploaded and analysed the TCR sequences and the gene expression matrices available on the 10X Genomics website. Utilising the available csv template in VDJView, we generated a third file containing the available metadata for each cell, e.g., subject ID, TotalSeq 15 surface markers including T cell differentiation markers (CD45RA, CD45RO, CCR7) and exhaustion and activation markers such as HLA-DR and PD-1, and tetramers read-counts (HLA-I restricted epitopes), MHC allele and other information. Given the large number of cells in the dataset, which can be a limitation for the standard computational resources available to the user, we used VDJView to randomly sample 15,000 cells from donor 1. We then performed quality control on the data, filtering out cells with >15% mitochondrial genes or abnormally high total expression counts, leaving 11,675 cells. After removing these obvious outliers, contaminants and poor quality cells, we filtered out cells with low tetramer read counts, or tetramer read counts that were not significantly higher than the negative control tetramers (also available in the dataset). This filtering resulted in 3,815 antigen specific T cells.

We used this set to explore the distribution of genes, markers for T cell differentiation, receptor clonotype, and tetramer specificity. Unsupervised analysis (Figure 2a) revealed 8 clusters with marker genes identifying signatures of cytotoxic activities of CMV, EBV and Influenza specific CD8+ T cells, and the presence of memory and naïve T cells (e.g., CCR7^+^ CD45RO^+^ and CCR7^+^ CD45RA^+^), thus, revealing clustering based on epitope specificity, T-cell differentiation and TCR specificity. Specifically, clusters 1 and 4 showed clonally expanded populations of EBV specific memory cells identified by marker genes being TCR V genes and by CDR3 specificity. Cluster 2 revealed influenza specific memory cells, expressing TRBV19, known to code for a public TCR specific to the highly-conserved M158-66 immunodominant epitope [21]. Clusters 3,5, and 6 mostly revealed CMV-specific cells displaying no obvious clonality. These three CMV-specific clusters revealed heterogeneous expression of Granzyme H and B genes, and of transcription factors LEF1, TCF7, and ZNF683 (Hobit), which are regulators of T-cell differentiation. The cells in cluster 6 expressed CCR7 and were clearly naïve (CD45RO^−^ CD45RA^+^). Finally, cluster 7 formed CMV and EBV specific and clonally expanded memory T cells, revealed by the same TCR CDR3 sequence. Notably, despite the filtering of low quality cells, cluster 8 revealed cells with reduced expression of all marker genes, including housekeeping genes RPL7 and RPL27, and with the highest percentage of mitochondrial genes, thus reinforcing the importance of quality control steps in scRNA-seq analysis.

**Figure 2.**
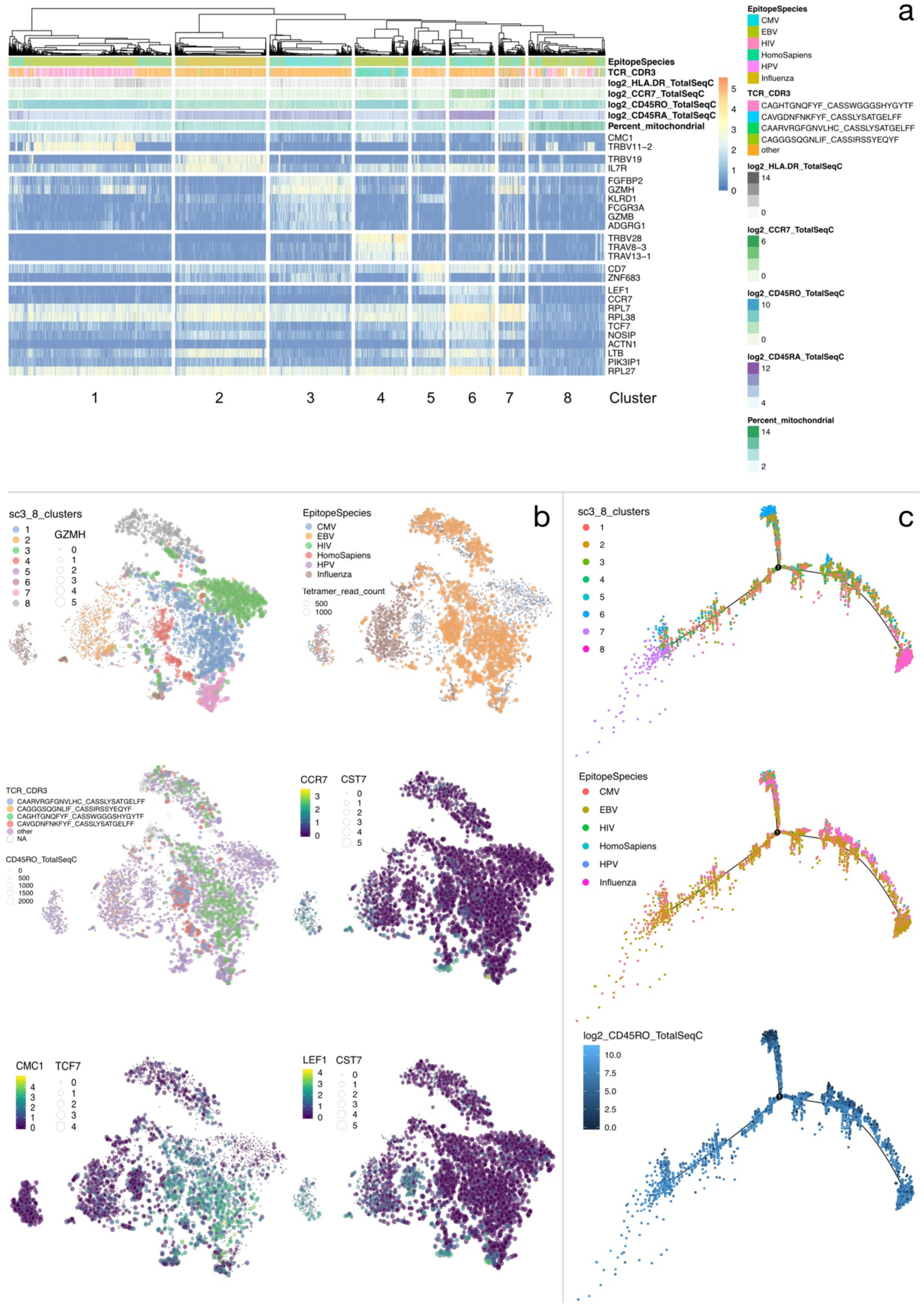
Analysis of CD8+ antigen-specific T cells sampled from Donor 1. **a**: Unsupervised clustering with k=8 clusters, p-value = 0.01, AUROC = 0.8. Epitope species specificity, the four largest TCR clones, surface protein expression levels, and the percentage of mitochondrial genes are annotated. **b**: t-SNE coloured by the results of clustering, epitope species, TCR clone and genes of interest (CCR7, CMC1, LEF1), with point size corresponding to highest tetramer read count of each cell, CD45RO TotalSeq expression, and genes of interest (GZMH, CST7, TCF7), show that clustering is preserved, and that clonally expanded T cells dominate the major clusters. Genes of interest reveal further sub-clusters of cells. **c**: Pseudo-time plots reveal a naïve to effector phenotype transition, with cluster preservation at the extremes of each state and a clear trajectory for influenza specific T cells.

We then utilised the dimensionality reduction features of VDJView to further explore clonality within these subsets. We used the t-SNE plots (Figure 2b) generated utilising the gene expression profiles to explore protein and tetramer expression, as well as other metadata information. As expected, the clusters identified via SC3 largely formed distinct clusters, with EBV and influenza specific T cells revealing the highest tetramer read counts, thus suggesting high affinity of these cells for the cognate antigens. Within the CMV and EBV specific T cells, clonally expanded T cells formed larger clusters, suggesting a common gene signature in clonally expanded populations. Interestingly, EBV and influenza tetramers, had higher read counts, suggesting that these cells might have higher binding affinities. By marking the expression of genes such as GZMH, LEF1, TCF7, CMC1 and CCR7 gene expression, the t-SNE plots revealed sub-clusters based on the differentiation status of T cells. Finally, we performed pseudo-time analysis (Figure 2c) to reveal a naïve to effector phenotype transition, shown by the increase in CD45RO expression, which is inversely mirrored in CD45RA expression. This analysis showed that naïve T cells identified in cluster 6 in the SC3 analysis formed a separate branch, while memory T cells were distributed across the pseudo-time structure.

We also analysed the TCRs of all T cells from donors 1 and 2. After performing the same quality control and filtering as outlined above, we were left with 55,922 antigen specific T cells (14,199 from donor 1 and 41,723 from donor 2). Both donors displayed clonally expanded populations (Figure 3), with 3 unique TCR expanded across at least 1,000 cells, and over 16 expanded across at least 100 cells. Both donors displayed VDJ gene usage bias, with a relatively high usage of TRBV19 common to both donors. We identified a total of 15,600 unique TCRs, with 411 TCRs common in both donors (Table 2 shows 15 of these). We also found evidence of cross reactive TCR that target different antigens within the same species, or across species, opening further avenues of study.

**Table 2:** TCR clones shared between donor 1 and donor 2, and the species they target with the number of occurrences in each donor.

| TCR_CDR3 | Species | d1 | d2 |
| --- | --- | --- | --- |
| CAGHTGNQFYF_CASSWGGGSHYGYTF | EBV | 2,379 | 37 |
| CAVGDNFNKFYF_CASSLYSATGELFF | EBV | 1,511 | 23 |
| CAARVRGFGNVLHC_CASSLYSATGELFF | EBV | 1,442 | 19 |
| CAASGYDYKLSF_CSVSASGGDEQYF | CMV, EBV | 214 | 10 |
| CAVFLYGNNRLAF_CSVSASGGDEQYF | CMV, EBV | 199 | 11 |
| CAASETSYDKVIF_CASSFSGNTGELFF | EBV | 38 | 5,810 |
| CADSGGGADGLTF_CASSLRDGSEAFF | EBV | 34 | 4,428 |
| CAASETSYDKVIF_CASSWGGGSHYGYTF | EBV | 14 | 35 |
| CAGAGSQGNLIF_CASSIRSSYEQYF | Influenza | 16 | 345 |
| CAVTDGGSQGNLIF_CASSIRSSYEQYF | Influenza | 120 | 39 |
| CAGAHGSSNTGKLIF_CASSIRSAYEQYF | Influenza | 71 | 48 |
| CAVSGSQGNLIF_CASSIRSSYEQYF | Influenza | 10 | 79 |
| CAAGGSQGNLIF_CASSIRSSYEQYF | Influenza | 10 | 77 |
| CAGGGSQGNLIF_CASSIRSSYEQYF | Influenza | 469 | 1,094 |
| CAGGGSQGNLIF_CASSVRSSYEQYF | Influenza | 119 | 72 |

**Figure 3:**
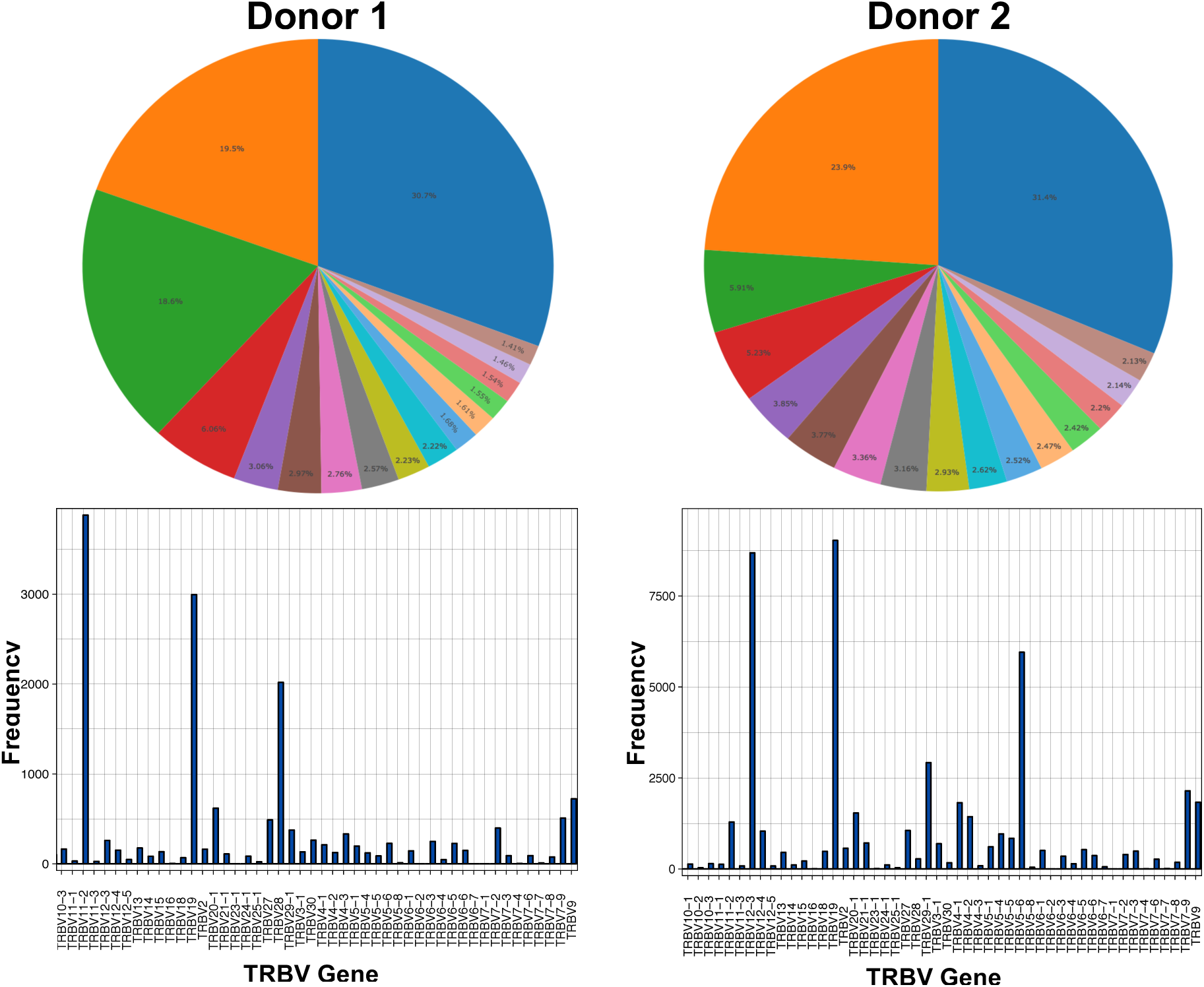
Summary of donor 1 and donor 2 clonal repertoires. Top 16 clones for each donor displayed in pie charts and the TRBV gene usage across all TCR in each donor is detailed in the histograms.

## Discussion

We have shown that integrating immune receptor and gene expression data with clinical information is useful to discover novel, biologically relevant findings from published data that do not emerge through previous analyses, and to further understand and discover medically relevant mechanisms. VDJView, a platform to conduct such analysis, forms an integrated set of known and novel tools that have a flexible design, expanding other tools and providing a robust quantitative framework to generate and study multi-omic immune cell data at the single cell level.

The proposed framework can be utilised by bioinformatics experts to develop and integrate new tools, as well as by clinical scientists and immunologists without profound knowledge of bioinformatics tools. Additionally, we propose that the software is a useful tool for lab-meetings as it promotes an on-the-go type of analysis that is suitable for quick hypothesis testing.

### Limitations

VDJView is developed in R, and therefore it is relatively simple to maintain and install. However, updates to the packages that VDJView utilises may cause dependency issues or loss of function due to code deprecation. This is an issue that requires periodic updates, and while we will maintain the software, fixing any issues that arise, we recommend using the suggested R versions. Additionally, while VDJView can analyse multiple datasets at once, the software does not perform any batch correction. As such, any technical variation in the data should be corrected for prior to uploading. VDJView will be maintained monthly and new tools will be integrated according to the development of software packages in the field, and the feedback received from users of the software.

## Conclusions

VDJView is a complete software package for downstream analysis of single cell gene expression, immune receptor and metadata, which allows exploratory and hypothesis driven analysis of multi-omic datasets. In summary, VDJView has the potential to allow clinical and experimental researchers to utilise complex genomics data to test biologically relevant questions.

## Availability and requirements

**Project name:** VDJView

**Project home page:** https://bitbucket.org/kirbyvisp/vdjview

**Operating system(s):** Linux, MacOS, with major features functional on Windows

**Programming language:** R

**Other requirements:** R 3.4.1 or higher

**License:** GNU

## Declarations

### Ethics approval and consent to participate

Not applicable

### Consent for publication

Not applicable

### Availability of data and materials

All data and metadata presented are publicly available and have been compiled into the following repository for ease of access: https://drive.google.com/drive/folders/1ATRZ239ubNu8Jv7Zy71WFylmGdC0TALt

### Competing interests

Not applicable

### Funding

FL acknowledges funding from NHMRC (APP1121643 project grant and Career Development fellowship APP1128416). JS and MG have PhD Scholarships from UNSW.

### Authors’ contributions

FL and JS designed the research. JS and SR wrote the code. JS and FL analysed the data and wrote the manuscript. All authors have read the manuscript and provided feedback.

## Acknowledgements

We thank Mandeep Singh, Katherine Jackson and Joanne Reed from the Garvan Institute of Medical Research, and Curtis Cai, Auda Eltahla, and Raymond Louie from the the Kirby Institute for Infection and Immunity for feature suggestions and use of the tool throughout its development.

## Notes

#### Summary of Updates

The revised paper solely documents VDJView, detailing added features, and presents analysis on a 10X dataset of over 150,000 CD8+ T cells with TotalSeq and dextramer staining.

https://bitbucket.org/kirbyvisp/vdjview

