## Supplementary Material for "Exploring and analysing immune single cell multi-omics data with VDJView"

### Supplementary Note 1: VDJView

#### VDJView input

##### Immune receptor data

Clonotype data obtained from software such as VDJPuzzle [1], the Cell Ranger TCR detection files from the 10X scRNA-seq 5' kit, BASIC [2], BraCeR [3], TraCeR [4] or similar tools can be uploaded to VDJView as comma-separated or tab-separated files. The files must be of MiGMAP (or similar) format, containing columns detailing the V, D and J genes, the CDR3 sequence of each receptor chain reconstructed. Additionally, VDJView accepts columns detailing contig expression, the isotype detected, whether the receptor is membrane bound or secreted, point mutations and overall mutation rates.

##### Gene expression matrix

Raw counts (non-normalized), FPKM, RPM, or TPM values obtained from gene expression quantification tools such as RSEM [5], Kallisto, Cufflinks or featureCounts can be uploaded into VDJView as a matrix or aggregate file. VDJView also accepts the matrix generated by Cell Ranger on 10X scRNA-seq data. VDJView accepts and merges multiple 10X Cell Ranger matrices, additionally, if a 10X Cell Ranger matrix file contains multiple matrices, VDJView will treat the first matrix as the gene expression data, and the second as additional metadata (this decision was made to load the recent 10X CD8 T cell data in which tetramer read counts and surface protein TotalSeq expressions were formatted into a second matrix within the same file as the gene expression counts). Cells in the gene expression data can be randomly sampled to reduce computational strain and resources required.

The clonotype and gene expression datasets are uploaded separately, allowing users to analyse them one at a time or simultaneously.

**Metadata**

Users can upload their own cell metadata as a csv file, each row detailing a cell (by ID) and its associated metadata. A CSV template of this file, tailored to the uploaded dataset, can also be downloaded.

#### VDJView features

**Quality Control (QC) and Normalisation**

- Gene expression data can be log normalised
- Violin plots show the distribution and relationships between common QC metrics; Number of genes expression, total expression counts, percentage of mitochondrial genes, and percentage of ribosomal genes
- Cells can be filtered in/out according to QC metrics using sliders

**Analysis**

- Ability to filter cells according to uploaded and application derived metadata (e.g. receptor type and CDR3 mutation rates)
- Tables summarising cell clonotypes, mutations, expression and isotype across individual chains and full receptor
- Pie charts showing clonotype usage across individual receptor chains and full receptor
- Histograms detailing V, D and J gene usage
- Histograms detailing CDR3 (amino acid) length distribution
- Circos plots detailing VDJ and chain pairing
- Scatter plots and tables to explore receptor repertoires shared between user-defined groups (tcR package)
- Dimensionality reduction (PCA, t-SNE and UMAP) plots. Plots support colour, size and shape specification based on uploaded metadata as well as perplexity selection for t-SNE. Plots also support point selection for further exploration of visual clusters.

- Unsupervised clustering performed by SC3 provides consensus matrices, gene expression and marker gene plots, with user specified number of clusters, P-value and AUROC and metadata display. The seed used for SC3 can also be specified to allow for reproducibility in otherwise stochastic plots
- Gene expression heatmaps for user-uploaded gene sets (performed by SC3)
- Once generated, SC3 clustering information is loaded into the metadata and can be visualised on other plots for more complex analysis of the unsupervised clustering results
- SC3 Clustering information can be downloaded in CSV format
- Supervised clustering performed by MAST providing gene expression plots and downloadable spreadsheets detailing gene fold change values
- Pseudo-time plots to determine single-cell trajectories run with monocle. Uploaded and application generated metadata, clustering results, and gene expression can be used to colour plot
- Most plots are customisable, and parameters can be changed mid-analysis

**Output**

- Generates downloadable metadata for isotype, mutation frequency, and membrane/secretion proteins for B cells
- Table displaying summary statistics for all uploaded cells and their associated metadata (cell metadata summary)
- All results/analyses are exportable into CSV files or high quality PDFs

**Installation**

- Works on both Linux and Mac OSX, with major features functional on Windows
- The software is self-contained and works on all recent R versions – the sole requirement is that R (3.4.1+) is downloaded
- Simple upload process with instructions on what files are expected

**Performance**

Running on a machine with 32GB RAM, a dataset of about 500 cells takes approximately 15 seconds to upload, most plots render instantaneously, with the exception the PCA, TSNE and UMAP plots which take about 10 seconds to render. Supervised/unsupervised clustering take 8/15 minutes to calculate, and the plots then take about 10 seconds to render. The Pseudo-time plot takes 10 minutes to calculate and then 10 seconds to render. The software has been tested on datasets of over 100,000 cells with gene expression data on over 40,000 cells, with the only limiting factor being computer memory.

### Supplementary Note 2: breast cancer scRNA-seq data

To demonstrate the utility and novelty of VDJPuzzle and VDJView, we analysed scRNAseq data from breast cancer tissue and associated metastatic lymph nodes from 11 subjects[6]. We utilized Chung et al., 2017 to show heterogeneous cell subsets and identification of T cells, B cells and macrophages using a priori gene signature sets and validation through experimental staining.

Here, we use VDJPuzzle for the identification of T and B cells as well as isotype detection, mutation estimation and membrane or secreted exon variant calling. Following this we feed the data generated by VDJPuzzle into VDJView. Of the total 563 samples, 14 bulk or pooled samples were removed, and we further analysed the remaining 549 samples. We identified 3 samples that carry both full-length BCR and TCR, thus suggesting that these samples were also multiple cells and not a single cell. We identified 103 B cells and 30 T cells using full-length receptor identification, and 180 cells with partial BCR (at least one chain) and 42 partial TCR. We identified 33 samples with partial BCR and TCR, thus suggesting that some of these cells may be the result of doublets or multiple cells per cell during sorting.

**Analysis of T and B cell clonotypes**

Excluding bulk and pooled samples, we found 138 IGH, 143 IGK and 125 IGL chains. In BC03 we found 69 IGH cells, of which 52 were from lymph node tissues. BC07 had 49 IGH chains, of which 42 were from lymph node tissues. As we report multiple isotypes for each cell it is not possible to estimate the exact combination of unique IGH-IGK/IGL chain pairing present in the cells, so instead, we report all possible combinations. We identified 178 TRA and 178 distinct TRB chains, and by excluding bulk samples we found 80 TRA and 40 TRB chains, with 38 complete αβ chains (**Supplementary Figure 1**).

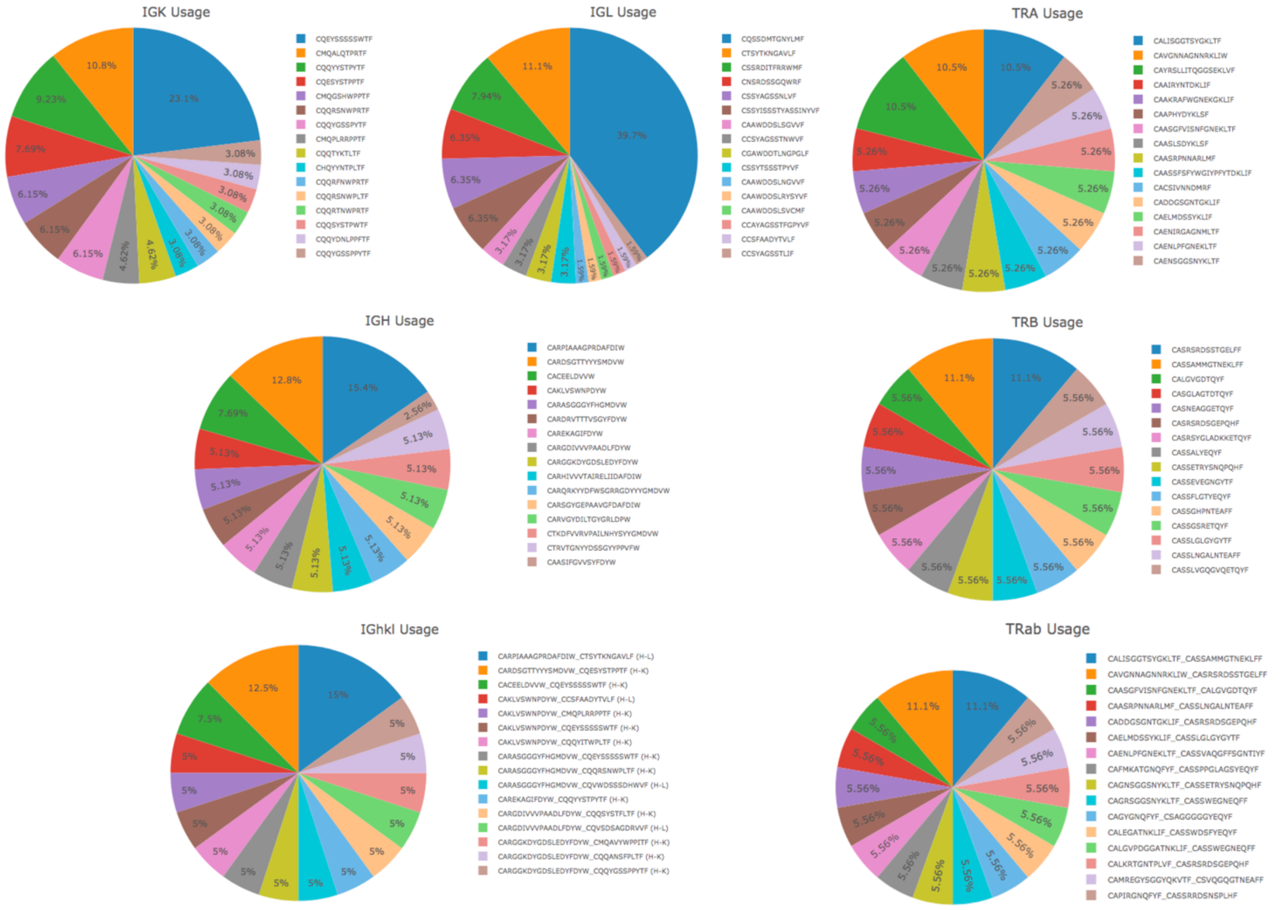

**Supplementary Figure 1**: T and B cell receptor CDR3 frequency for Chung et al[6] immune cells (only the 16 most abundant CDR3 contigs are shown per chain).

#### Analysis of shared clonotypes and receptor encoding V, D, and J genes

The analysis of these immune cells revealed a highly diverse clonal repertoire (**Supplementary Figure 1**). The TCRαβ clone identified in subject BC03 (CAVGNNAGNNRKLIW-CASRSRDSSTGELFF, with germline TRAV8-3, TRAJ38, TRBV10-2 TRBD2, TRBJ2-2) was found in one cell from primary and one from metastatic tissues. We identified 31 clones across the 101 full-length BCRs, with shared clonotypes between primary and metastatic tissues and also between subjects, suggesting that a common repertoire of B cells is involved in breast cancer with a migratory pattern between primary and metastatic tissues. B cells from the lymph nodes were more clonally diverse than those found in breast cancer, with 27 clones in 64 cells from the lymph node and 7 in 37 cells from the breast tissue. Notably, there was a significant overlap in gene usage between B cells across tissue types (Supplementary Table 2.9); for instance, IgHV4−59 was expressed in 6 B cells from breast cancer tissues, and 10 lymph node derived cells. This analysis also revealed a full-length BCR clone (CDR3 CACEELDVVW-CQEYSSSSSWTF, IgHV3-33, D2-15, J1, and IgKV1-

5, J1) was found in 2 cells of BC09 breast tissue and 1 cell in metastatic tissue of BC03. This K chain (V1-5, J1) was expressed in 15 cells across subjects BC04, BC07, BC09 and BC03LN.

We observed that many BCRs were shared across cells taken from different tissues as well as from different subjects. These clones are summarised in **Supplementary Table 1** below. Of note are CDR3 clones CARPIAAAGPRDAFDIW-CTSYTKNGAVLF and CACEELDVVW-CQEYSSSSSWTF which are expressed in cells from both the primary breast cancer tissue, as well as the metastatic lymph node. Additionally, the IgK CDR3 of this second clone, was present in 8 cells taken from the breast cancer tissue of subjects BC07, BC04 and BC09 as well as in 7 cells taken from the lymph node of subject BC03. Using a similar method in VDJView, we explored the V D and J genes of the BCR present across cells derived from each tissue as well as from each subject. The genes are summarised in **Supplementary Table 2**. Despite the large number of different genes (especially IgH V genes), a similar gene pool was used to encode the BCRs across tissues, explaining the inter-tissue clonality observed in **Supplementary Table 1**. This can be seen in **Supplementary Figure 2** which shows the IgH V genes used in the BCRs of cells derived from the primary (a) and metastatic (b) tissues. Additional CDR3 clones were identified by analysis of the CDR3 IgL chain, e.g. CQSSDMTGNYLMF (IgLV3-25/IgLV3-16, J2), which is present in all 25 cells with a BCR from subject BC02.

**Supplementary Table 1**: Common single chain and full-length BCR CDR3s in cells across multiple subjects grouped by tissue of origin. Subjects BC01, BC05, BC08 and BC11 have few immune cells and have been omitted from the table.

| **CDR3** | **Primary Breast Cancer** | | | | | | | **Metastatic Breast Cancer** | |
|---|---|---|---|---|---|---|---|---|---|
| | BC03 | BC07 | BC04 | BC06 | BC09 | BC10 | BC02 | BC03LN | BC07LN |
| (IgH) CARPIAAAGPRDAFDIW | | | | | 4 | 2 | | | |
| (IgK) CMQALQTPRTF | | | | 5 | | | | | |
| (IgK) CQESYSTPPTF | | | | | | | | 5 | |
| (IgK) CQEYSSSSSWTF | | 1 | 1 | | 6 | | | 7 | |
| (IgK) CQQYYSTPYTF | 3 | | 1 | | | | | | 2 |
| (IgL) CNSRDSSGQWRF | 2 | 1 | 1 | | | | | | |
| (IgL) CQSSDMTGNYLMF | | | | | | | 25 | | |
| (IGL) CTSYTKNGAVLF | | | | | 5 | 2 | | | |
| (IgH_IgL) CARPIAAAGPRDAFDIW-CTSYTKNGAVLF | | | | | 4 | 2 | | | |

| | | | | | | | | |
|---|---|---|---|---|---|---|---|---|
| (IgH_IgK) CACEELDVVW-CQEYSSSSSWTF | | | | | 2 | | | 1 |

**Supplementary Table 2**: Common V and J genes expressed in BCR heavy (IGH), and light (IGK and IGL) chains across multiple subjects grouped by tissue of origin. Subjects BC01, BC05, BC08 and BC11 have few immune cells and have been omitted from the table.

| **Gene** | **Primary Breast Cancer** | | | | | | | **Metastatic Breast Cancer** | |
|---|---|---|---|---|---|---|---|---|---|
| | BC03 | BC07 | BC04 | BC06 | BC09 | BC10 | BC02 | BC03LN | BC07LN |
| IgHJ3 | 1 | | | | 5 | 2 | | 2 | 10 |
| IgHJ4 | 9 | 4 | 1 | | 2 | | | 22 | 19 |
| IgHJ5 | 3 | 1 | | | | | | 6 | 5 |
| IgHJ6 | 4 | 1 | | 6 | | | | 20 | 5 |
| IgHV1-18 | 1 | 1 | | 1 | | | | 5 | 2 |
| IgHV1-46 | | | | 2 | 1 | | | 1 | 1 |
| IgHV1-69 | 1 | | | | | | | 1 | 5 |
| IgHV3-23 | | 1 | | | | | | 4 | 2 |
| IgHV3-30 | | 1 | | 1 | | | | 2 | 3 |
| IgHV3-30-3 | 2 | | | | | | | 6 | |
| IgHV3-33 | 1 | | | | 2 | | | 3 | 1 |
| IgHV3-48 | 1 | | | 1 | 1 | | | | 2 |
| IgHV4-39 | 3 | 1 | | | | | | 4 | 1 |
| IgHV4-59 | 1 | 1 | | 1 | 5 | 2 | | 4 | 2 |
| IgKJ1 | 6 | 2 | 2 | 5 | 6 | | | 27 | 8 |
| IgKJ2 | 9 | 1 | 2 | 1 | 1 | | | 8 | 9 |
| IgKJ3 | 2 | 2 | | | 1 | | | 12 | 4 |
| IgKJ4 | 3 | 4 | | 2 | | | | 6 | 10 |
| IgKJ5 | 1 | 1 | | | | | | | 4 |
| IgKV1-39 | 1 | 2 | | | 1 | | | 9 | 7 |
| IgKV1-5 | 2 | 1 | 2 | | 6 | | | 10 | 2 |
| IgKV1-9 | | 1 | | | | | | 1 | 2 |
| IgKV2-28 | 1 | 1 | | 5 | | | | 6 | 1 |
| IgKV2-30 | 4 | | | | | | | 1 | 1 |
| IgKV3-11 | 2 | | | | | | | 12 | 3 |
| IgKV3-15 | 2 | | 1 | | | | | 3 | 1 |
| IgKV3-20 | 1 | 1 | | 1 | | | | 6 | 2 |
| IgKV4-1 | 5 | 2 | 1 | | 1 | | | 2 | 4 |
| IgLJ1 | 3 | | | 1 | | | | 7 | 2 |
| IgLJ2 | 9 | 6 | 4 | 4 | 11 | 2 | 25 | 22 | 26 |
| IgLV2-14 | 1 | 1 | | 1 | 5 | 2 | | 7 | 4 |

| | | | | | | | | | |
|---|---|---|---|---|---|---|---|---|---|
| IgLV2-23 | 1 | 1 | | | | | | 2 | 3 |
| IgLV3-25 | 3 | | 1 | | | | 16 | 6 | 2 |

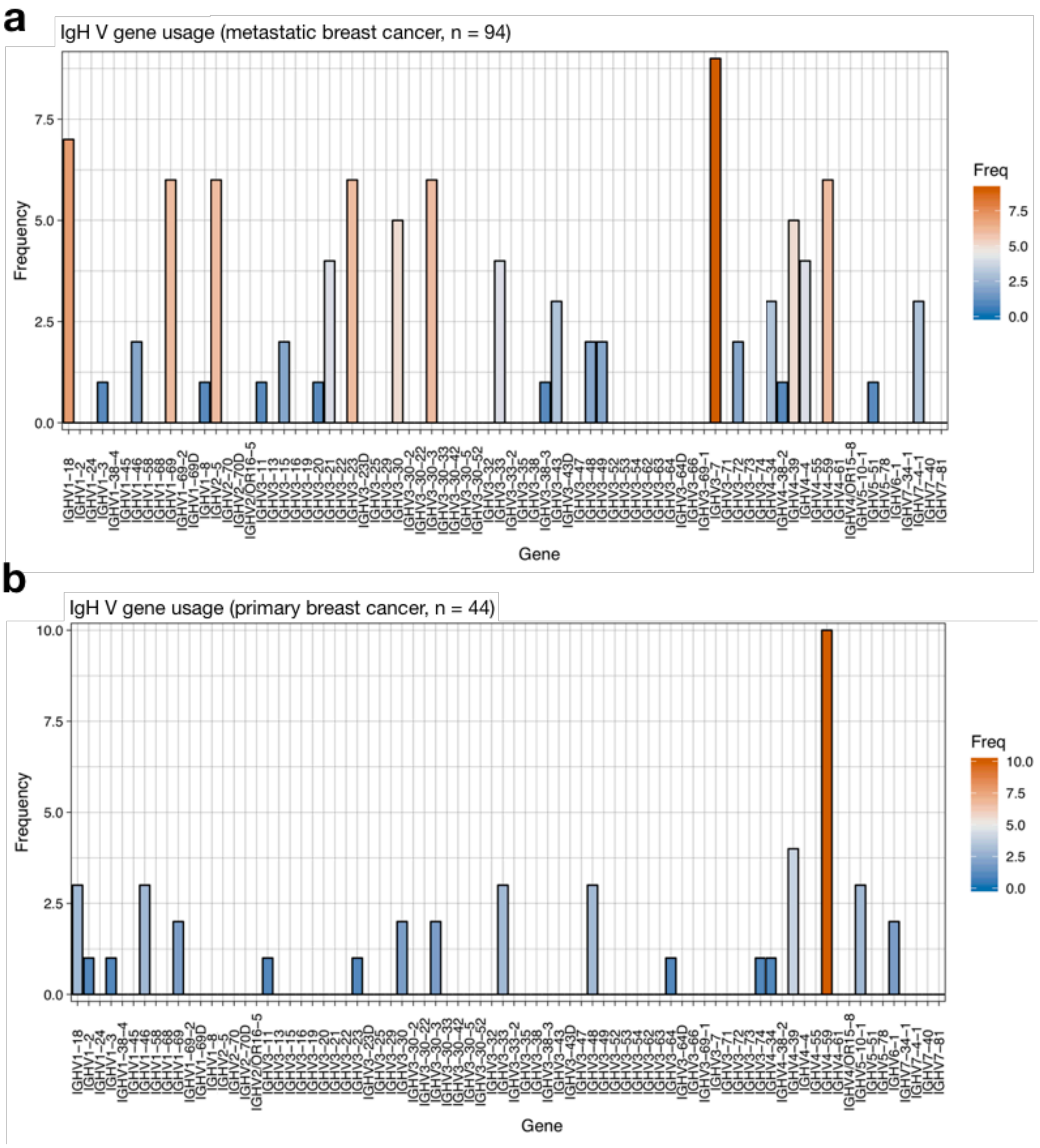

**Supplementary Figure 2**: IgH V gene usage across cancer immune cells with a reconstructed IGH BCR chain. a) Immune cells from metastatic breast cancer tissue, b) Immune cells from primary breast cancer tissue.

#### Dimensionality reduction and integration of TCR and BCR sequences

We have also re-analysed the data using three dimensionality reduction tools (PCA, tSNE and UMAP), all of which are available on VDJView. We showed that the three tools provide similar results. Notably, by exploring the relationship between the available metadata and clonotype data, we found that B and T cells from the double positive (ER+ and HER2+) tissues had a gene signature overlapping with the B cells derived from the triple-negative and were present in two major clusters, mostly separated from the other cell types (**Supplementary Figure 3**). Dimensionality reduction performed on the B and T cells only revealed a cluster of lymph node derived B cells in double positive tissues, while again both T and B cells had a similar gene signature (**Supplementary Figure 3**).

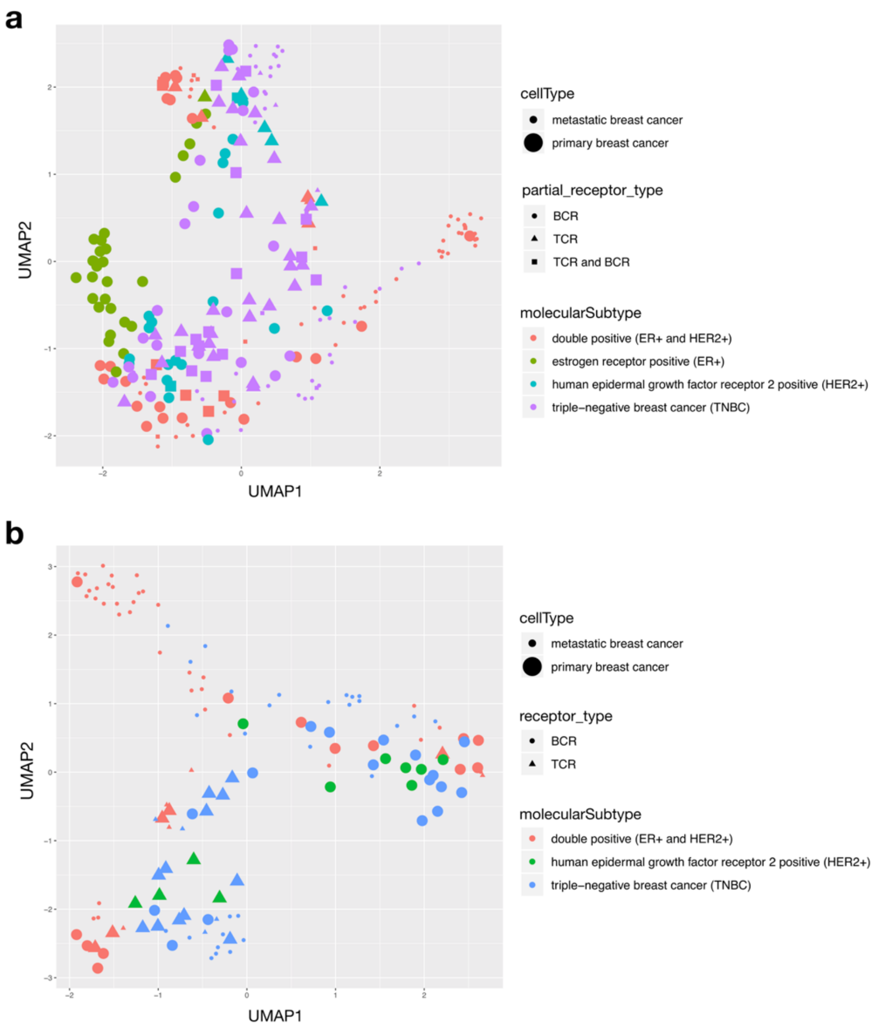

**Supplementary Figure 3**: UMAP dimensionality reduction of cancer single cells, with colour, size and shape determined by molecular subtype, tissue of origin and receptor type respectively. a) All immune cells with a partial or full-length receptor, b) Cells with either a full-length TCR or BCR.

**Unsupervised analysis of immune cells identified with VDJPuzzle**

To further investigate the heterogeneity of T and B cells, we used VDJView to integrate immune receptor information with the gene expression profile and available metadata, and performed unsupervised clustering as well as a model-based differential expression analyses focussing on comparing B and T cells between tissues as well as between cancer molecular subtypes. **Supplementary Figure 4** shows the result of running unsupervised clustering with SC3 through VDJView on all 245 cells that carried a partial or full-length receptor, which revealed evidence of 8 clusters and clearly demonstrating that immune cells form distinct clusters based on identity (B and T cells), isotype, tissue of origin and molecular subtype. In this analysis, cells with a partial BCR clustered with those with a full-length BCR, and cells with a partially reconstructed TCR clustered with cells with a full-length TCR, thus suggesting that those partial reconstructions are effectively real immune cells and not contaminated samples. T cells largely formed a single cluster with marker gene CD96 associated to immune modulation, as well as expression of IL2R-γ and FYB which is known to control IL-2 secretion. The rest of the clusters were largely composed of B cells based on tissue of origin, molecular subtype of cancer, and notably a cluster that was composed of somatically hypermutated IgG1 B cells in metastatic lymph-node of double positive breast cancer. The majority of these cells also expressed a secreted form of the heavy chain, and a gene signature suggesting a highly active and differentiated B cells expressing LRMP, TCL1A, RGS13, PAX5, HMGN2P5, HMGB2, CD22, and EAF2. These cells are likely plasmablast following a reactivation of memory B cells. In this cluster, the over-expression of PAX5 and TCL1A could also indicate presence of malignant immune cells as these genes are often found in leukemia and likely to contribute to oncogenesis BCL6 [7, 8]. A second cluster of membrane-bound B cells was identified in primary breast tissue of patient BC02, with high expression of MGP, SERPINA and ESR1. Interestingly, this cluster contained only cells with a partially reconstructed BCR.

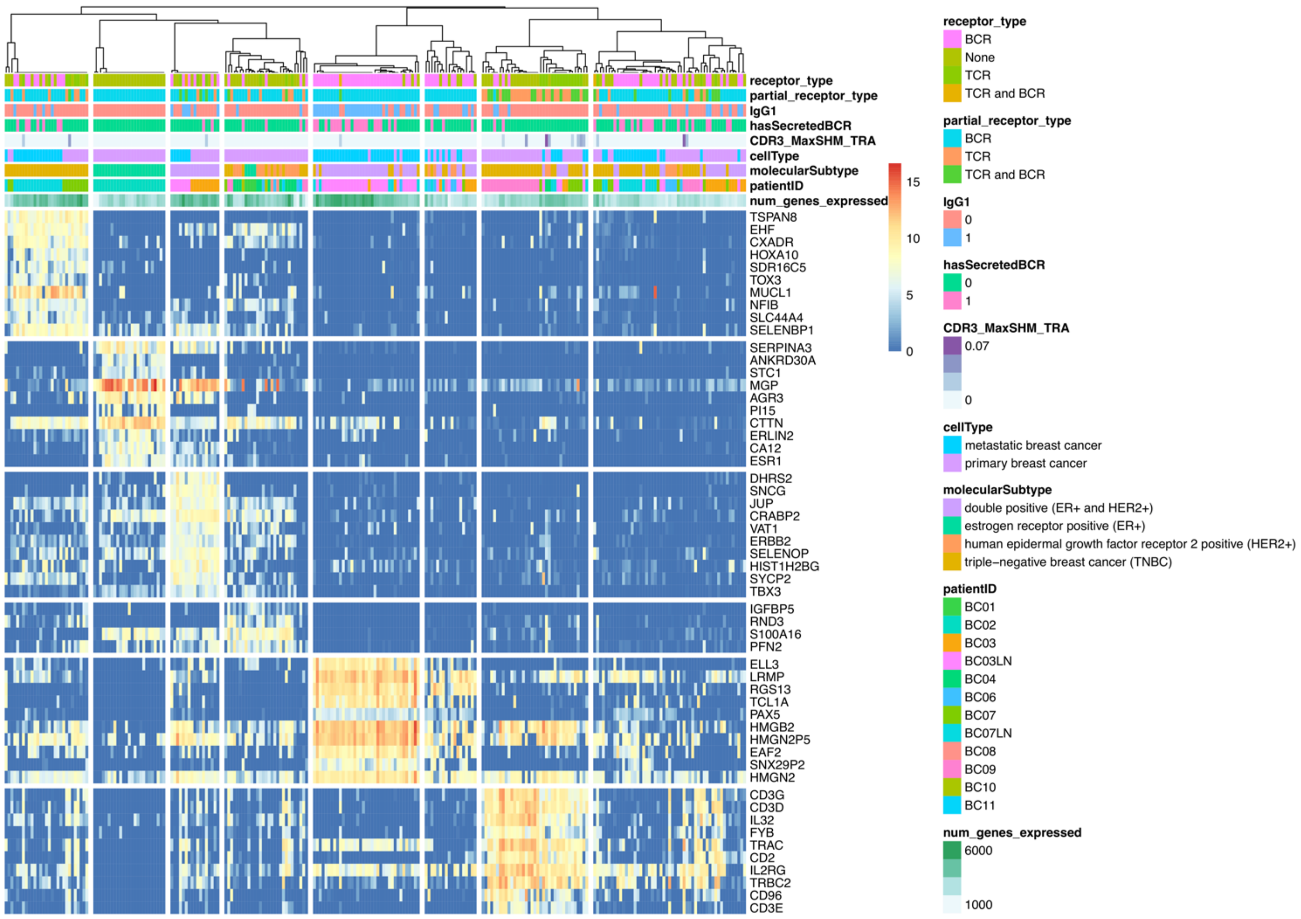

**Supplementary Figure 4**: Unsupervised clustering using SC3 from scRNA-seq of all the T and B cells identified from VDJPuzzle with full-length or single chain receptors. Marker genes show clear separation between T and B cells, and with greater heterogeneity among B cells. B cell clusters are largely identified by patient, cancer molecular subtype, and tissue or origin (primary and metastatic tissues). Notably a large cluster of IgG1$^+$ active B cells and with secreted form of BCR are identified within lymph-node with elevated SHM and marker genes associated to B cell differentiation within germinal centre (ELL3, EAF2, PAX5).

To validate the findings presented above, we used the clustering output of SC3 and investigated the expression profiles of immune subsets utilising a list of known genes associated with B and T cell phenotypes (**Supplementary Figure 5**). This analysis showed that B cells residing in the lymph node were IgG1$^+$, with gene signature CD24$^-$, CD22$^+$, CD38$^+$, and CD69$^+$. In contrast, other cells, mostly in the breast cancer and some in metastatic tissues, were CD24$^+$, CD22$^-$ and CD38$^-$ . Clusters were also separated by the expression of CXCR4, BCL6, and CD40. All together this profile suggests that highly active B cells exist in the lymph node and express several genes that are associated to support cytotoxic T cell response such as.

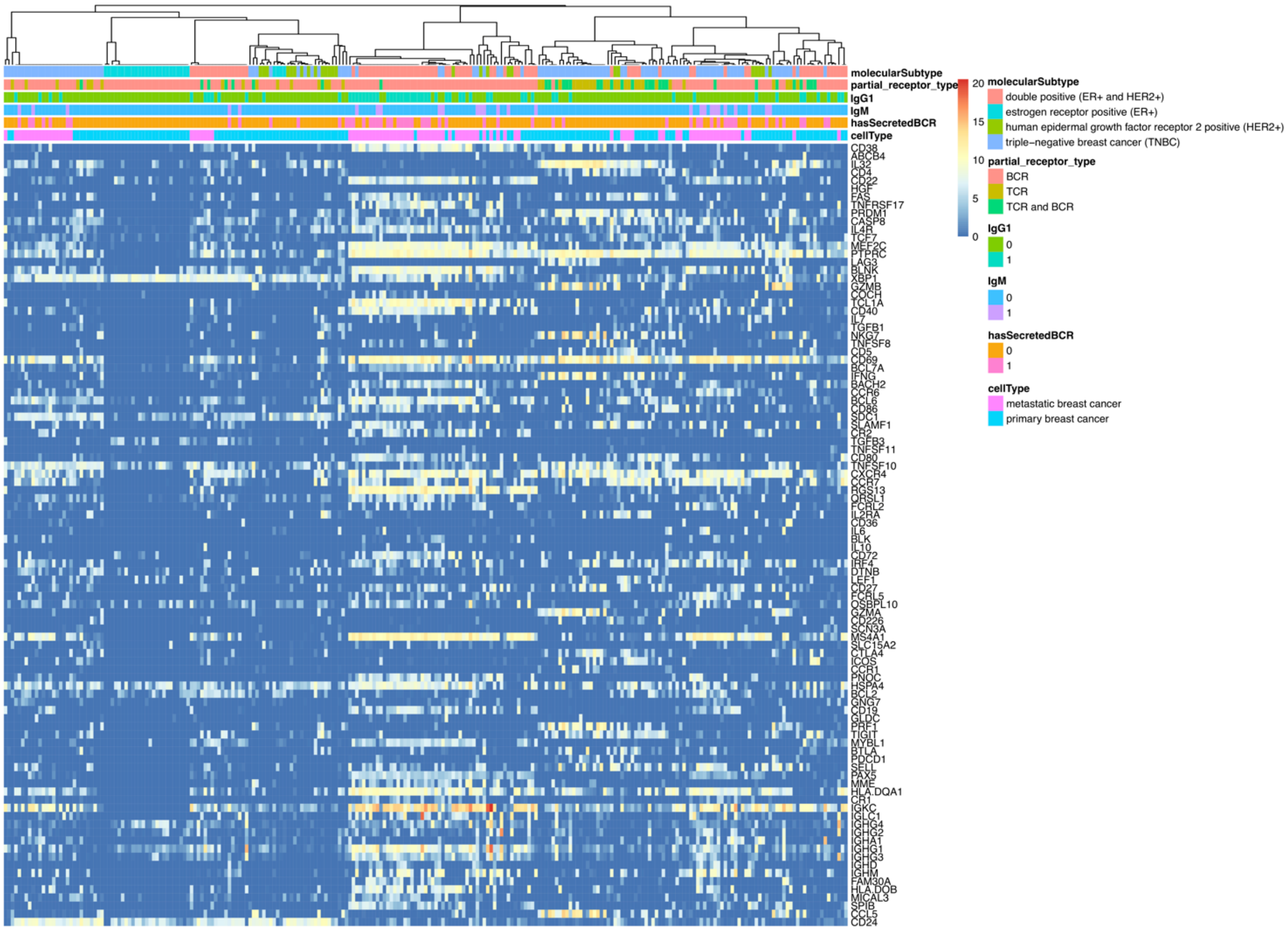

**Supplementary Figure 5:** Unsupervised gene expression analysis of T and B cells using gene list of known immune cell phenotype specific genes, confirming that cells cluster together according to the tissue of origin (cell type), as well as the receptor type and B cell isotype.

#### Comparison of immune cells using a model-based approach

To further elucidate the molecular signatures of these immune cells, we performed a model-based differential expression and clustering analyses utilizing MAST in VDJView. Specifically, we analysed gene profiles of immune cells by molecular subtypes of breast cancer (**Supplementary Figure 6**) and by tissue of origin (primary and metastatic, **Supplementary Figure 7**). Differential expression analysis comparing T and B cells between double positive (ER+ and HER2+) and triple negative breast cancers revealed distinct clusters of B cells by isotype and SHM levels. We discovered B cells found in metastatic lymph nodes from the subject BC03 affected by double positive breast cancer tissues were consistent with plasmablast cells, as these cells expressed a secreted form of IgG1, elevated SHM, and expression of genes such as MZB1 and EAF2, and ELL3. This cluster was also confirmed in the analysis comparing immune cells between tissue of origin. In contrast, B cells found in the primary tissue in subjects with triple negative breast cancer were mostly resting naïve or memory cells, with IgG1, IgG3 and IgM isotypes, expressing HLA DR markers, as well as CD99[9], thus implying that they function as antigen presenting cells.

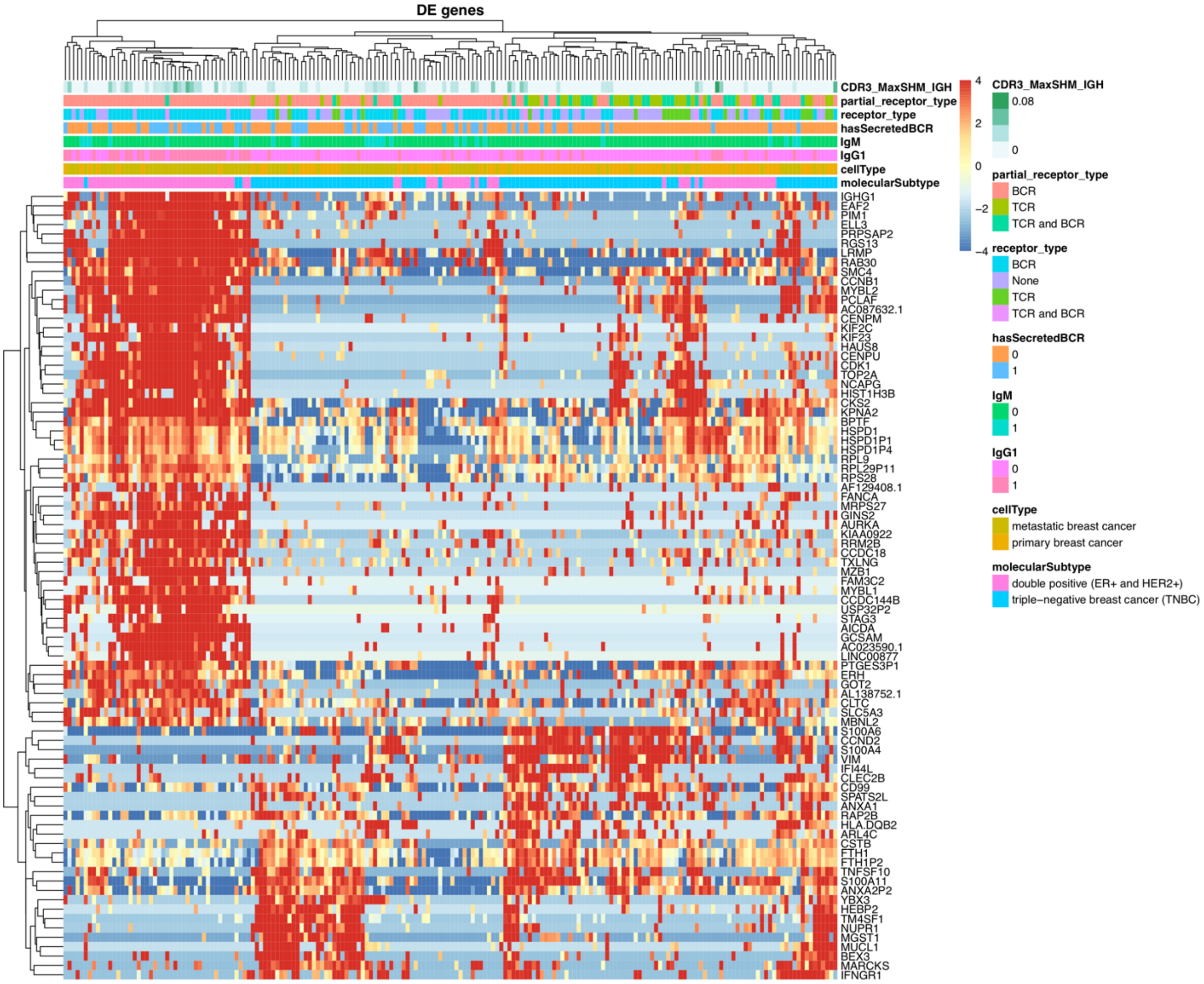

**Supplementary Figure 6**: Heatmap of immune cells clustered by cancer molecular subtypes. Supervised clustering depicting differentially expressed genes (fold change > 2) between ER+HER2+ and triple-negative breast cancer subtypes. The cells in the ER+HER2+ cluster (left) are primarily IgG1+ cells with an elevated SHM, secreted BCR from the metastatic lymph node tissue, and have gene expression profiles consistent with plasmablast B cells.

A MAST differential gene expression analysis was performed to compare the immune cells in primary and metastatic tissues. In the resulting heatmap, **Supplementary Figure 7**, we see that cells in the first cluster (on left) had a high expression of marker genes such as antigen presentation associated markers (HLA DR, CD79), CD53 (associated to activation of memory B cells), CD69 as well as markers of B cell differentiation (MS4A1 which encodes CD20). The cells with IgG1 (compared to a subset of IgM in this cluster) had a higher number of genes expressed, and an increased expression of genes such as D1PR2 and PAX5, and activation markers within germinal centre (CD83,

FCRLA, CD22 and EAF2, which mediate apoptosis in Germinal centre). New genes were identified such as TCF4, and GPR18B. Notably, this cluster also appeared to be enriched in genes that have been found in lymphoma and hyperplasia (TCL1A, CD79B, IRF4 and IRF8).

Cells that formed the second cluster (in middle) were derived from breast cancer tissues, and were mostly partially reconstructed B cells with a gene expression profile enriched for genes encoding for the S100 protein, Cyclin D1, MGP and ANXA1, a membrane-localized protein that binds phospholipids which has been associated with breast cancer. The final cluster (on right) was formed by B and T cells. The B cells were IgG1 and IgM, characterised by the expression of CD53, TCF4, as well as CD83 and HLA-DR, thus again suggesting a role in antigen presentation.

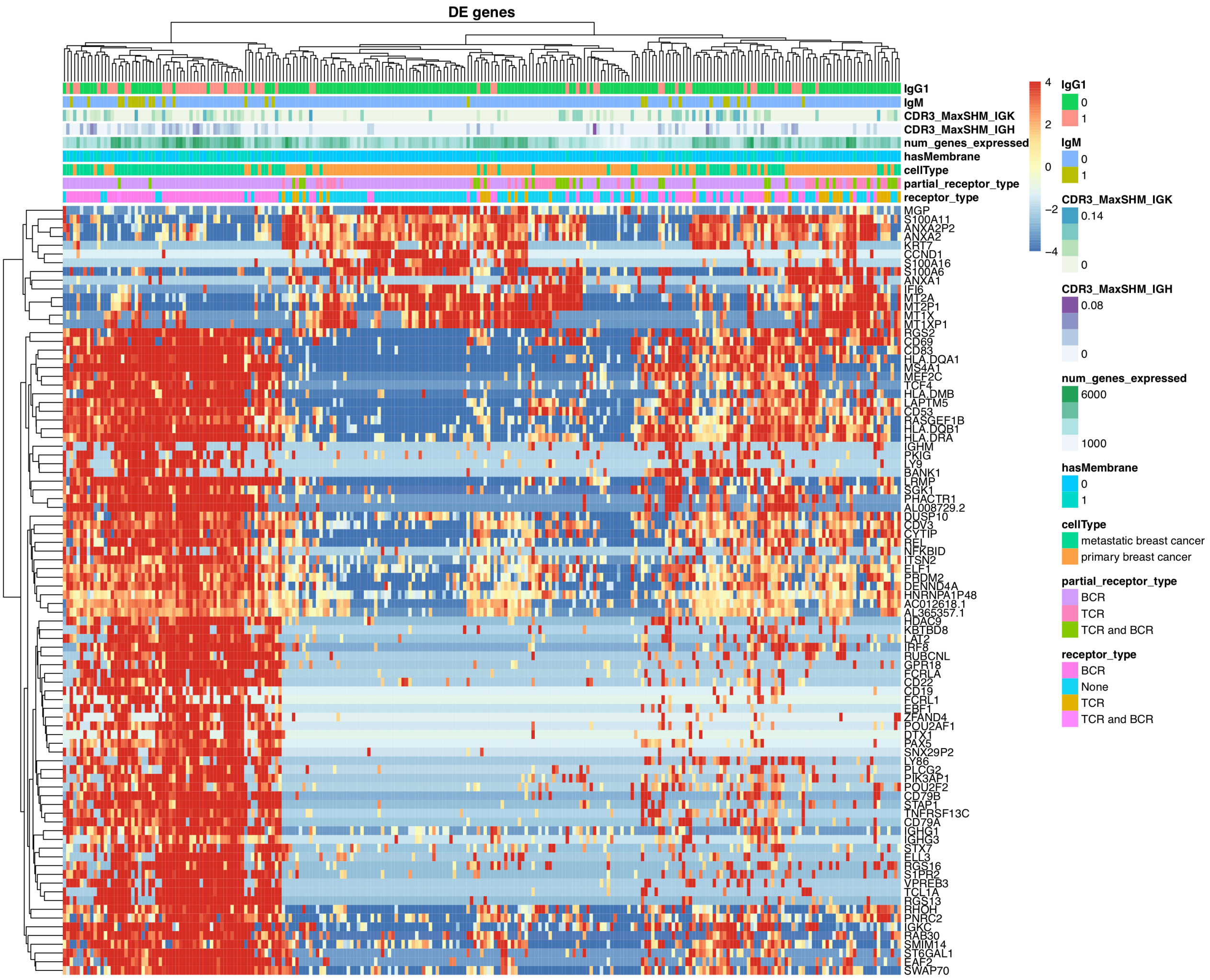

**Supplementary Figure 7**: Heatmap of fully and partially reconstructed immune cells clustered by tissue of origin (cellType). Supervised clustering depicting marker genes (fold change > 2) between primary and metastatic (lymphoid) tissues of origin. The cells in the metastatic lymph node cluster (left) are primarily IgG1+ B cells, while primary breast cancer cells lacking a full TCR or BCR cluster together (middle).
